# A dynamic and combinatorial histone code drives malaria parasite asexual and sexual development

**DOI:** 10.1101/2021.07.19.452879

**Authors:** Hilde von Grüning, Mariel Coradin, Mariel R. Mendoza, Janette Reader, Simone Sidoli, Benjamin A. Garcia, Lyn-Marie Birkholtz

## Abstract

A ‘histone code’ defines system-level crosstalk between histone post-translational modifications (PTMs) to induce specific biological outcomes. Proteome-scale information of co-existing PTM across the entire chromatin landscape of the malaria parasite, *Plasmodium falciparum,* was lacking. Here, we used advanced quantitative middle-down proteomics to identify combinations of PTMs in both the proliferative, asexual stages and transmissible, sexual gametocyte stages of *P. falciparum*. We provide an updated, high-resolution compendium of 72 PTMs on H3 and H3.3, of which 30 are novel to the parasite. Co-existing PTMs with unique stage distinction was identified, indicating a dynamic and complex histone code with increased connectivity of novel PTMs seen in gametocytes. Chromatin proteomics of a gametocyte-specific combination, H3R17me2K18acK23ac, identified a SAGA-like effector complex (including the transcription factor AP2-G2) tied to this combination to regulate gene expression in mature gametocytes. Ultimately, this study unveils previously undiscovered histone PTMs and their functional relationship with co-existing partners. These results highlight that investigating chromatin regulation in the parasite using single histone PTM assays might overlook higher order gene regulation for distinct proliferation and differentiation processes.

## Introduction

Histone N-terminal tails are reversibly modified by an array of covalent histone post-translational modifications (PTMs). These alter chromatin structure to fine tune gene expression in most eukaryotes, resulting in changes in cell fate. Although the contribution of individual histone PTMs in particular biological processes are well described (Berger, 2002; Zhao Y. & Garcia, 2015), there is growing evidence to indicate that histone PTMs not only function individually, but also act in a concerted manner to direct transcriptional programmes according to the cell’s immediate needs. The outcome of such coordination ultimately define a cell’s fate and function (Turner, 2000), such as to proliferate (Klein *et al*., 2019; Schwammle *et al*., 2016), differentiate (Bhanu *et al*., 2016; Chen T. & Dent, 2014), or become quiescent (Liu *et al*., 2013; Young C.P. *et al*., 2017). This association between histone PTMs that work in coordination has been postulated to constitute a functionally-relevant, unique pattern or ‘histone code’ (Jenuwein & Allis, 2001; Strahl & Allis, 2000).

As example of the importance of histone PTM crosstalk, the presence of histone H3, serine 10 phosphorylation (H3S10ph) impairs the binding of the effector protein heterochromatin protein 1 (HP1) to the well-known repressive PTM, H3K9me3 (Hirota *et al*., 2005). This blocks cellular differentiation in mouse embryonic stem cells (Fischle *et al*., 2005; Johansen & Johansen, 2006). Such drastic changes in gene regulation and cellular fate can also be effected by combinations of PTMs that include typically less abundant PTMs e.g. H3K9ac and H3K14ac affect H3R8me2 (Fulton *et al*., 2018; Kirmizis *et al*., 2007).

The histone PTM landscape in the causative agent of severe malaria in humans, *Plasmodium falciparum*, is associated with a dynamic abundance profile of individual PTMs that changes during various developmental stages of the parasite (Coetzee *et al*., 2017) and contributes to a tightly controlled transcriptional programme (Hollin & Le Roch, 2020). Several of the individual histone PTMs are essential to various biological processes, as demonstrated by gene knockout or chemical perturbation of histone ‘writer’ and ‘eraser’ enzymes responsible for histone PTM placement and removal, respectively, detrimental to parasite development (Coetzee *et al*., 2020; Zhang *et al*., 2018).

The parasite undergoes rapid rounds of cell division during its asexual replication to proliferate every ∼48 h. A small percentage of asexual parasites differentiate to gametocytes through five distinct stages of development (stages I-V) over ∼14 days in *P. falciparum*, after which mature male and female stage V gametocytes can be transmitted to the mosquito vector (Josling *et al*., 2018; Maier *et al*., 2019). The parasite’s chromatin organisation fluctuates between mostly euchromatic in asexual parasites, characterised by transcriptionally permissive PTMs of H3K9ac and H3K4me3 (Bartfai *et al*., 2010; Bozdech *et al*., 2003; Salcedo-Amaya *et al*., 2009), and more heterochromatic states during gametocytogenesis, marked with H3K9me3, H4K20me3 and extended HP1 occupancy (Coetzee *et al*., 2017; Flueck *et al*., 2009). However, nuanced distinctions exist between the different developmental stages during gametocytogenesis and underscores the transcriptional differences between the early- (stage II/III) and late-stage (stage IV/V) gametocytes (van Biljon *et al*., 2019). Typical heterochromatic PTMs such as H3K27me3 and H3K36me3 are exclusive to the immature, early stages (Coetzee *et al*., 2017; Connacher *et al*., 2021) whilst more mature stages do contain euchromatic PTMs (H3K4me3 and H3K79me3) in preparation for onwards transmission and gamete formation (Coetzee *et al*., 2017).

Evidence of histone PTM combinations in *P. falciparum* is sparse but includes methylation of H3K4 or H3K9 by the histone lysine N-methyltransferase SET7, only in the presence of already acetylated H3K14 (Chen P.B. *et al*., 2016). H4K8ac, as a likely regulator of parasite proliferation in asexual parasites (Gupta *et al*., 2017), is also found in combination with H4K5ac, H4K12ac and H4K16ac as a result of the acetyltransferase activity of MYST (Miao *et al*., 2010). Quantitative chromatin proteomics alluded to the presence of additional co-existing histone PTMs in *P. falciparum* parasites, with prominent stage specificity and increased presence during gametocytogenesis (Coetzee *et al*., 2017; Saraf *et al*., 2016).

The majority of co-existing histone PTM pairs are typically identified with a peptide-centric proteomics pipeline, frequently named “bottom-up”. Histones are digested with trypsin or other enzymes that generate relatively short peptides prior MS analysis. With this workflow, co-occurrences between histone PTMs can be identified only for those which localize nearby in the amino acid sequence. However, this proteomic approach is not suitable for distal co-occurring PTMs. As such, distinctively modified peptides and co-occurring PTMs cannot be accurately identified and quantified (Janssen *et al*., 2019). The development of “middle-down” MS has advanced proteomics and allowed the complexity of PTM combinations to be investigated (Sidoli & Garcia, 2017). Although technically challenging, middle-down MS allows evaluation of longer histone tails of ∼50-60 residues (Sidoli *et al*., 2016a; Sidoli *et al*., 2017) to accurately and simultaneously identify and quantify hundreds of combinatorial PTMs (Moradian *et al*., 2014; Sidoli *et al*., 2017). With this powerful approach, system-level crosstalk of multiple, interacting co-existing PTMs defined clear, combinatorial histone codes in nematodes (Sidoli *et al*., 2016b), mammalian cells (Schwammle *et al*., 2016; Sidoli *et al*., 2017; Tvardovskiy *et al*., 2017; Tvardovskiy *et al*., 2015), cells undergoing epithelial to mesenchymal transition (Garabedian *et al*., 2018; Jiang T. *et al*., 2018; Schräder *et al*., 2018; Sidoli *et al*., 2017; Sweredoski *et al*., 2017) and stem cell reprogramming (Benevento *et al*., 2015).

Here, we present the systems-level identification and characterisation of the combinatorial histone code of *P. falciparum* parasites, which was generated using quantitative, middle-down proteomics. We identify a comprehensive histone code that is dynamic and has distinct fingerprints in different life cycle stages, implying refined functions to allow different biological outcomes associated with parasite pathology and survival. Several co-existing PTMs are involved in direct crosstalk, indicative of coordinated function, particularly in gametocytes, with the code in immature gametocytes being most connected and mature gametocytes using unique combinations. This is exemplified by the functional association of H3K18acK23ac that interacts with a unique reader complex in mature gametocytes to enable strategy-specific gene expression. With this first, comprehensive report of a combinatorial histone code in a eukaryotic parasite, we show that *P. falciparum* relies on dynamic interactions between histone PTMs for development and differentiation. This data could serve as a model for the importance of combinatorial histone PTMs in pathogenesis in protista.

## Results

### Adapting middle-down proteomics for P. falciparum parasites

To investigate the presence of co-existing histone PTMs with middle-down proteomics, we isolated parasites as enriched trophozoites (92 ± 0.9%), immature early-stage gametocytes (83 ± 3.1% stage III gametocytes, 14 ± 2.2% stage II), and mature gametocytes (94 ± 2.9% stage V), to allow stage-specific differences to be inferred (Fig 1A). All samples yielded > 300 000 cells / sample, resulting in sufficient yields of acid-soluble nuclear protein fractions containing the histones: 17 ± 1.1 μg for the trophozoite population and 58 ± 10 μg and 70 ± 16 μg for the immature and mature gametocytes, respectively (Fig EV1A).

**Figure 1.**
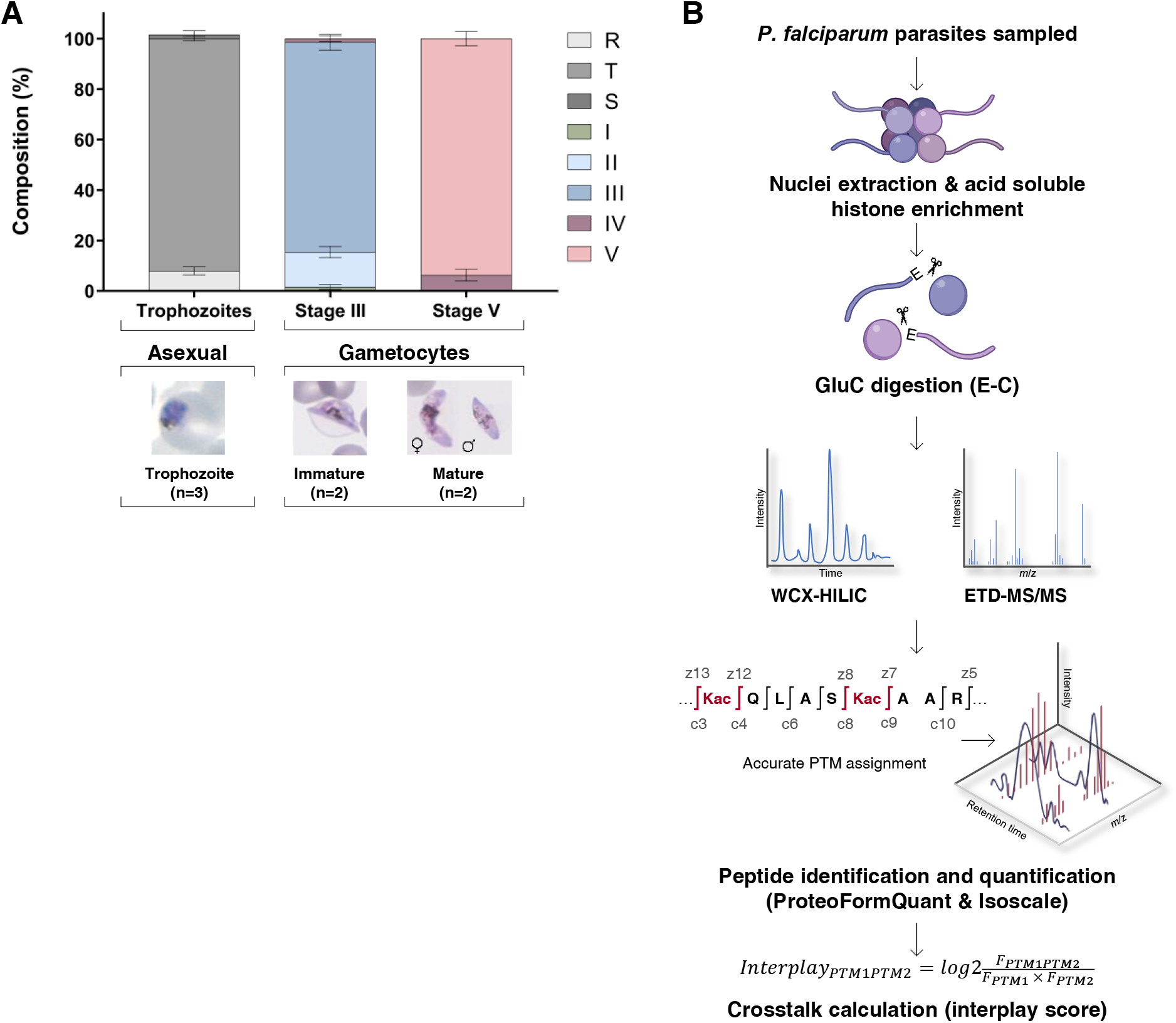
Middle-down MS workflow for analysis of *P. falciparum* parasite. **(A)** The stage composition of the three-biological stage analysed in this study and representative morphology. Trophozoites (mean ± SEM) samples contained a small percentage of ring (R) and schizont (S) stages, while the stage III samples (mean ± SD) contained stage I (I), II (II), III (III) and IV (IV), while stage V samples (mean ± SD) consisted of stage III, IV and V gametocytes. The trophozoites were from three independent biological repeats and gametocyte stages from two independent biological repeats. **(B)** The middle-down proteomics workflow. Histones were enriched from trophozoite, stage III and stage V gametocytes and digested using endoprotease GluC which cleaves at the C-terminal of glutamic acid residues, generating an intact N-terminal histone H3 peptides (50 amino acid residues in length). To allow sample loading in aqueous buffer and for the most efficient separation for histone N-terminal tails, nano liquid chromatography equipped with a two-column system consisting with a C18-AQ trap column and a weak cation exchange-hydrophilic interaction chromatography (WCX-HILIC) resin analytical column coupled online with high resolution tandem mass spectrometry (MS-MS) fragmentation was performed using electron transfer dissociation (ETD). Spectra were identified using the Mascot and peptides were quantified using isoScale. The flask image was adapted from the image from Servier Medical Art (http://smart.servier.com/). Servier Medical Art is licensed under a Creative Commons Attribution 3.0 License (CC BY 3.0 license: https://creativecommons.org/ licenses/by/3.0/). Data processing and analysis workflow involved MS spectral deconvolution using Xtract (Thermo Fisher Scientific) followed by database searching using Mascot (Matrix Science, UK) with files generated from PlasmoDB (https://plasmodb.org/) and subsequent removal of ambiguously mapped PTMs and stringent quantification (including co-fragmented isobaric species) using IsoScale Slim (http://middle-down.github.io/Software). The relative abundance of an individual PTM (PTM1/2) is calculated by summing the relative abundances of all proteoforms carrying the specific individual PTM. The interplay between two individual modifications is calculated by dividing the observed abundance of bivalent PTMs (FPTM1PTM2) with the predicted frequency of a combinatorial PTM (FPTM1 x FPTM2). FPTM1PTM2 is calculated by summing the relative abundances of all proteoforms carrying both PTMs.

An optimised middle-down proteomics workflow was established to accurately identify and quantify individual and combinatorial histone PTMs at scale (Fig 1B)(Coradin *et al*., 2020; Holt *et al*., 2019). The native histones from each sample were digested with GluC endoproteinase to produce polypeptide fragments of ∼50-60 amino acids in length (≥5 kDa) that were separated and identified by high-resolution nano liquid chromatography-MS/MS; data processing and peak extraction was performed with in-house developed tools, ProteoFormQuant and HistoneCoder (Greer *et al*., 2018; Jung *et al*., 2013). Unique PTMs (with a false discovery rate [FDR] <1%) with sufficient fragment ions (Fig EV1B) were accurately quantified, with a coefficient of variance ≤33% and average Pearson r^2^=0.69 between biological replicates (Fig EV1C), allowing comparison of PTM quantities between samples. To ensure differentiation of isobaric peptides as a result of the middle-down proteomics, data-dependent acquisition data were further processed using isoScale Slim (Sidoli *et al*., 2017), that extracts the total fragment ion intensity of histone tail spectra as representative of their abundance, and it discriminates isobaric forms using unique site-specific fragment ions using the principle of the fragment ion relative ratio (FIRR) (Pesavento *et al*., 2006). From the accurately identified and quantified histone PTMs per peptide, co-existing frequency was calculated for the observed presence of combinatorial PTMs (from MS2 level evidence) as a function of predicted co-existence frequencies, defined as an ‘interplay score’ (Sidoli *et al*., 2014).

### A high-resolution, quantitative compendium of histone PTMs in Plasmodium from middle-down proteomics

The high-resolution, sensitive nature of middle-down proteomics allowed successful identification and quantification of 83 PTMs on histone H2B.Z, H3 and H3.3 (Fig 2), as only these histones have GluC cut sites on their N-terminal histone tail (Coradin *et al*., 2020). Mass tolerance was set at 30 ppm to accurately distinguish between lysine acetylation (42.011 Da) and lysine trimethylation (42.047 Da) as previously determined (Sidoli *et al*., 2014). 72 of the 83 identified PTMs were confidently quantified; the others were only detected but with a signal insufficient to assess an abundance (Fig 2). As no enrichment was performed for phosphorylations, these could not be quantified. A few PTMs previously identified with H2B.Z (K14ac, T30ac, K42ac), H3 (S10ac, S10ph, S28ph, S32ph) and H3.3 (S10ac, T11ac, S22ac, S28ph, S32ph) could not be confirmed (Coetzee *et al*., 2017; Saraf *et al*., 2016). Since these PTMs were only previously qualitatively described, their presence remains to be confirmed.

**Figure 2.**
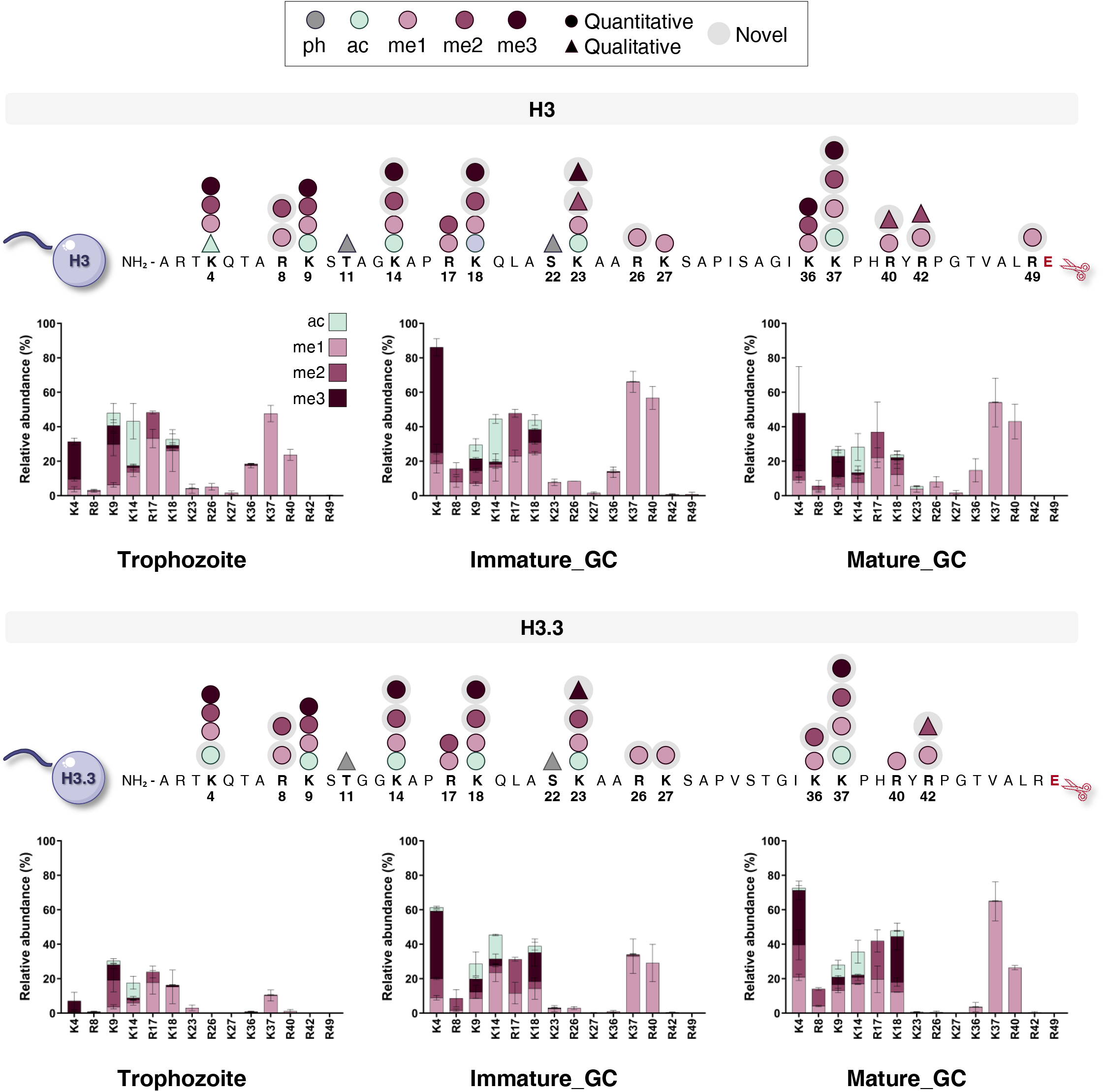
The relative abundances of individual PTMs on histone H3 and variant histone H3.3 from *P. falciparum* trophozoite, immature and mature gametocytes. The histone PTM landscape as shown for histone H3, H3.3 and H2B.Z (Fig EV2). A total of 83 PTMs on histone H3, variant histones H3.3 and H2B.Z were identified across all stages, including 72 quantitative (circle) and 12 qualitative histone PTMs (triangle) of which 35 were novel and detected for the first time in *P. falciparum* parasites (grey shaded). Abbreviations denote the PTM which included mainly acetylation (ac) and mono-, di- and trimethylation methylation (me1, me2 and me3, respectively) and phosphorylation (ph). The N-terminal peptide fragmented for analysis is shown with corresponding amino acid sequence. The relative abundances of the individual histone PTMs on histone H3 and histone variant H3.3, showing relative abundances on the different histone positions with either acetylated (teal), monomethylation (me1, light purple), dimethylation (me2, medium purple) and trimethylation (me3, dark purple). Data are from >2 independent biological repeats, mean ± SEM.

Of the 83 PTMs identified (Fig 2), 35 were described for the first time in *P. falciparum* parasites, of which 30 could be accurately quantified. As expected, the rarely observed variant H2B had the least number of PTMs. This included six known PTMs but also one novel PTM (H2B.ZK18me3, Fig EV2). Histone H3 contained 40 PTMs of which 33 was accurately quantified, including 13 novel PTMs. These include acetylation and methylation of H3K37, and methylation of several arginines (H3R8me1/2, H3R26me1, H3R40me2, H3R42me1 and H3R49me1). Several PTMs displayed high relative abundances (>20%) in more than two stages (e.g. H3K4me3, H3K14ac, H3R17me1, H3K18me1, H3K37me1 and H3R40me1), with the novel PTM, H3K37me1, highly abundant in immature gametocytes (70 ± 6%) (Fig 2). These changes in abundance levels between stages is also evident for H3K4me3, H3R17me2 and H3R40me1, which increased significantly from trophozoites to immature gametocytes (*P*≤0.01, n=≥2, Fig EV3). Histone H3.3 contained 37 PTMs (33 quantified) with substantially higher abundances of PTMs modified in both gametocyte stages compared to asexual parasites than seen for H3. Methylation PTMs were again abundant including H3.3K4me1/3, H3.3K18me3, H3.3R17me2 and H3.3R40me1, with H3.3K37me3 significantly increased abundance in mature gametocytes (*P*≤0.01, n=2, Fig EV3).

Collectively, we demonstrate that middle-down MS identified >80 unique PTMs and quantified 88% thereof. This includes 30 novel modifications, thereby providing an updated, high-resolution compendium of histone modifications across multiple life cycle stages of *P. falciparum*.

### A stage-specific combinatorial histone PTM code exists in P. falciparum parasites

The middle-down proteomics dataset was used to identify and quantify co-existing PTMs on H3 and H3.3 across the three life cycle stages of *P. falciparum*. On average, three co-existing PTMs were present on any given peptide for both H3 and H3.3 (Fig 3A), similar to what is seen for these two histones in humans and mice (Garcia *et al*., 2007; Tweedie-Cullen *et al*., 2012; Young N.L. *et al*., 2009). H3.3 is the exception, with on average only two co-existing PTMs present in trophozoites but as many as seven (e.g. H3K4me1R8me1K9me1K14acR17me1K18acK37me1) in immature gametocytes (Appendix 1).

**Figure 3.**
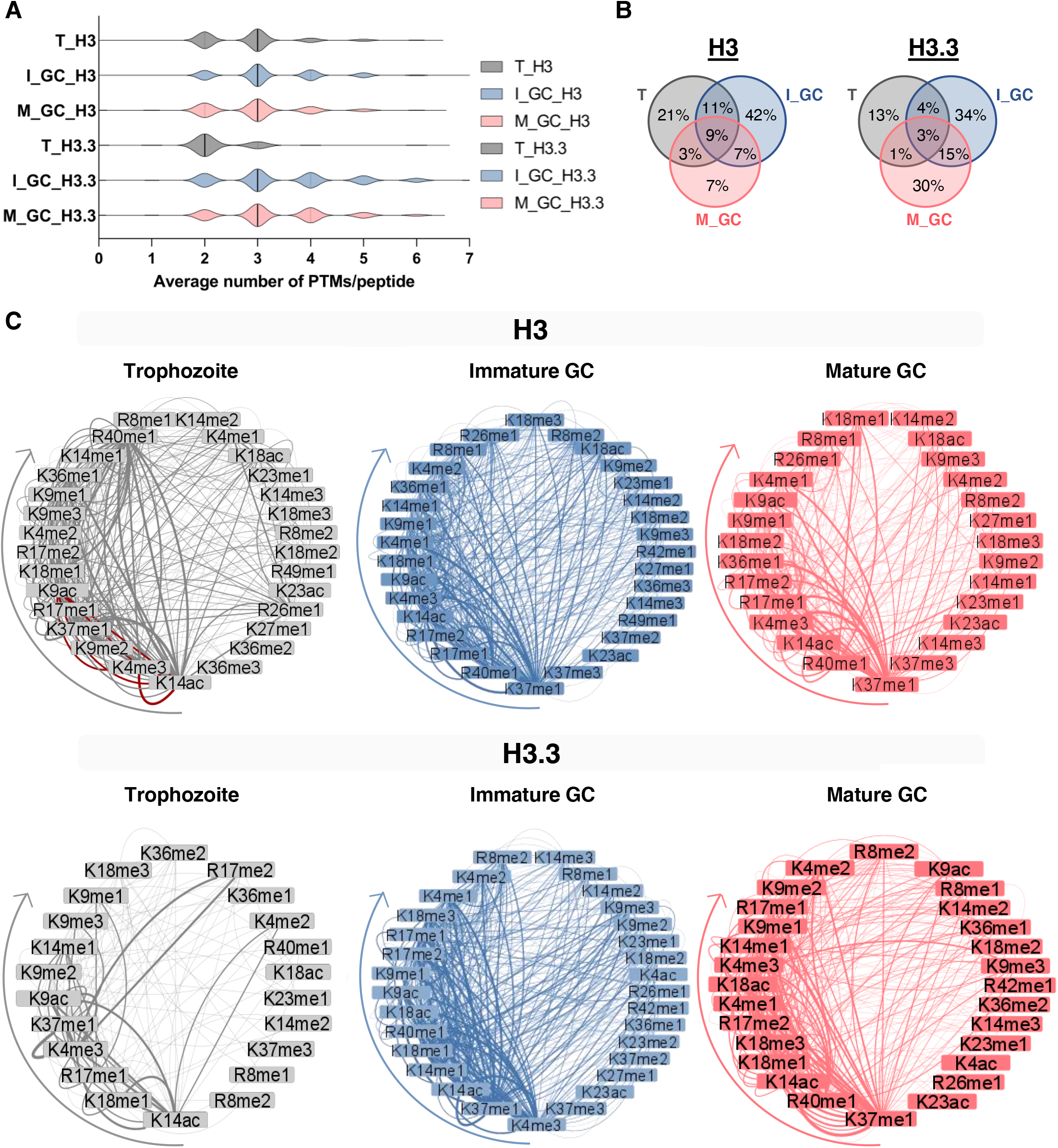
Prominent combinatorial histone PTM reorganization in *P. falciparum* parasites and gametocytes. **(A)** Violin plots show the distribution of the number of PTMs that are present on histone H3 and variant H3 tails where the dotted line indicated the upper and lower quartiles and the solid line indicating the median. The lower limit represents an unmodified histone tail. T: Trophozoite; SIII: Stage III and SV: Stage V gametocyte. **(B)** The Venn diagram indicates combinatorial peptides shared between the three stages from histone H3 and H3.3 were relatively unique to each stage (trophozoites, grey; stage III, blue; stage V, pink) sharing only 9 % and 3 % of combinatorial peptides, respectively. **(C)** The co-existing histone PTMs observed on histone H3 and H3.3 for trophozoites, stage III and stage V gametocytes are visualised as ring plots where the nodes are the histone PTMs while the edges represent the connection to another co-existing partner PTM. All combinations are included in the supplementary data. The arrows represent the start of the most connected PTM (left) toward least connected (right) in a clockwise direction. The co-existence of H3K9ac with H3K4me3, H3K14ac with H3K4me3 and H3K9ac with H3K14ac are highlighted with red edges in the trophozoite stage. The arrows indicate the most prevalent PTM to the least prevalent PTM in co-existence.

Stage-stratification was clearly evident in the histone PTM combinations, with immature gametocytes displaying the highest proportion of unique combinations (42%) on H3, with only a minor proportion (9%) of combinations shared between asexual parasites, immature and mature gametocytes (Fig 3B). Stage-specific diversity was somewhat less evident for H3.3 (Fig 3B), with a markedly decreased number of co-existing PTMs present in trophozoites for this histone (Fig 3C). The stage-stratification was characterised by changes in the identity of the most prevalent combinations. Trophozoites are characterised by a large number of combinations involving H3K14ac, H3K4me3, H3K9me2, and novel PTMs H3K37me1 and H3R17me1. These include the well-characterised, known combination of the archetypical euchromatic PTMs H3K4me3 and H3K9ac (Cui & Miao, 2010; Salcedo-Amaya *et al*., 2009), with H3K14ac due to the coordinated action of SET7 (Chen P.B. *et al*., 2016) and the histone acetyltransferase GCN5 (Fan *et al*., 2004a, b). In gametocytes, the novel PTMs H3K37me1 and H3R40me1 are involved in the highest number of co-existing PTMs, with further specification seen between immature gametocytes (with frequent interactions with H3R17me1&2 present) and mature gametocytes (higher connectivity for H3K14ac and H3K4me3 than H3R17me1). Combinations involving H3R42me1 occurred exclusively in gametocytes. Arginine methylation may therefore well make up a key component of the histone code in *P. falciparum* parasites along with other histone PTMs, particularly for gametocyte stages.

This data indicates that histones are rarely modified by single PTMs. Most frequently, co-existing modifications dictate chromatin fine tuning. As well, these histones codes are re-arranged in position and type of modifications in different life cycle stages of the malaria parasite. The most frequent combinations in gametocytes diverge from those trophozoites, with immature gametocytes associated with the highest number of co-existing PTMs ascribed to the presence of novel PTMs. The parasite therefore employs a unique and diverse set of PTM combinations likely to guide stage-specific gene expression.

### Histone PTM pairs display unique crosstalk to assert function

The extent and relevance of the influence between pairs of co-existing PTMs was subsequently interrogated by quantifying the co-existing frequency as an interplay score (IS) for bivalent combinations (Appendix 2). This provides a metric to predict the likelihood of individual pairs of histone PTMs to either: 1) have a positive interplay score to indicate co-dependence on one another, suggesting that one PTM require the presence of another PTM to exert its biological function (Kirsch *et al*., 2020; Schwammle *et al*., 2014; Sidoli *et al*., 2014); or 2) have a negative interplay score to indicate mutual exclusivity and/or functional independence (Hunter, 2007)(Fig 4). If two PTMs are randomly deposited on the chromatin in an independent manner, they will have an interplay score close to zero.

**Figure 4.**
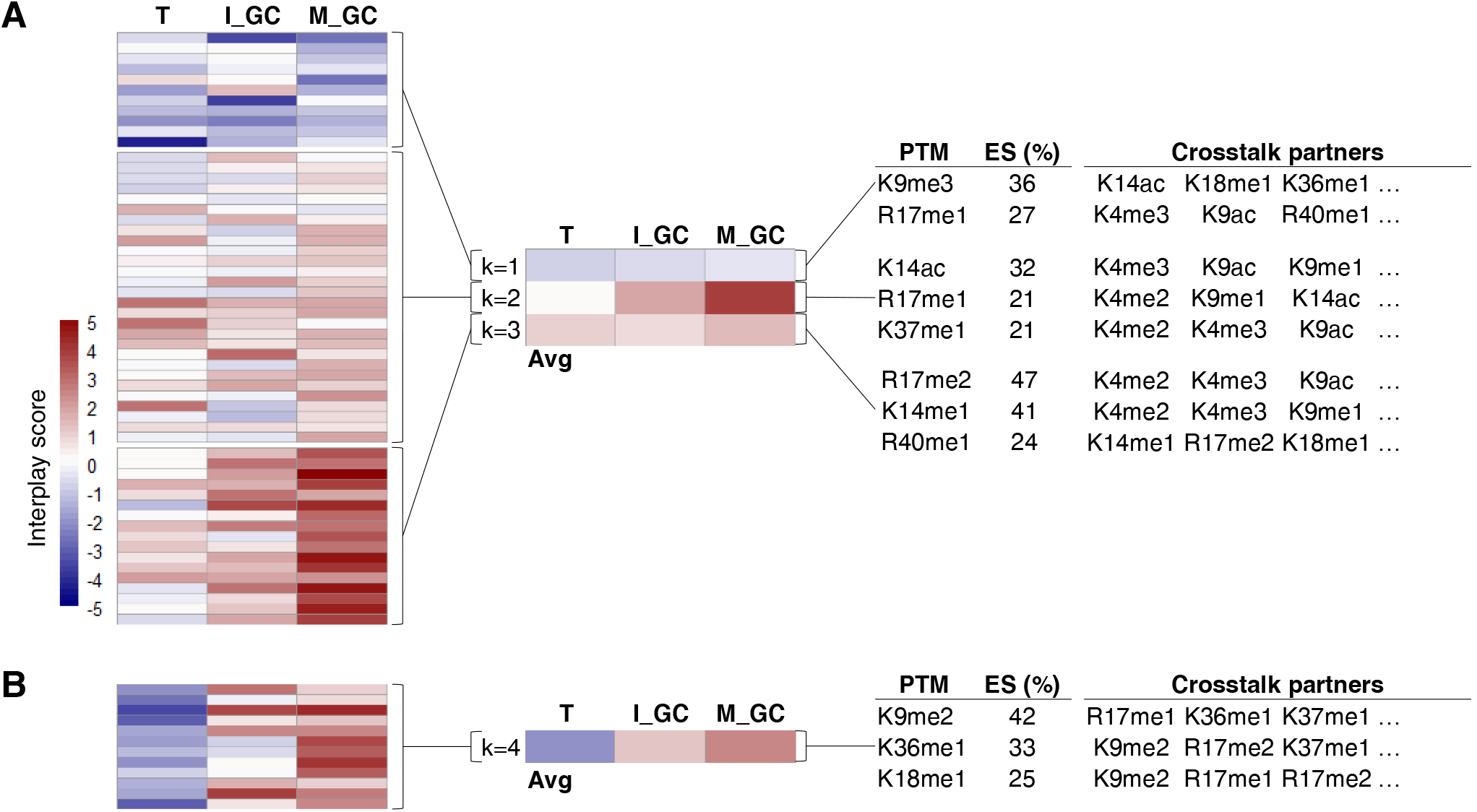
Histone modification crosstalk conserved across life cycle stages in *P. falciparum*. **(A)** Interplay scores for PTM combinations consistently present in all life cycle stages analysed. Heatmaps shows k-means clustering (complete, k=4) of the overlapping 35 % or 68 bivalent combinations and the respective interplay scores shared between all three stages. The average interplay scores are summarised for each cluster. **(B)** Interplay scores in cluster 4 across the life cycle stages, and PTMs that show enrichment within these clusters are quantified with an enrichment score. An enrichment score percentage (ES %) is shown for the top PTMs that are over-represented in each cluster. The enrichment score is calculated by dividing the total number of times a given PTM is present by the total number of combinations in the cluster. Selected crosstalk partner PTMs are indicted, with all the combinations provided in Fig EV4.

A third of all co-existing histone PTMs had similar connectivity and co-existing frequency profiles between all three stages of development of *P. falciparum* (Fig 4). Of these, 11 combinations show negative interplay scores throughout development (Fig 4A), which includes multiple combinations of the repressive PTM H3K9me3. For example, this PTM is found on the same histone molecule with H3K14ac significantly more rarely than stochastic co-occurrence in all three stages, implying mutual exclusion and thus opposing biological functions. The typically euchromatic signal of H3K14ac is therefore negated in the event of H3K9me3, and this contributes to the HP1-bound heterochromatic state as described for certain gene sets (Brancucci *et al*., 2014; Flueck *et al*., 2009; Perez-Toledo *et al*., 2009). This combination is found also in differentiated stem cells (Gonzales-Cope *et al*., 2016) where H3K9me3 act as a barrier to cell reprogramming induced in pluripotent stem cells, regardless of the presence of the activating H3K14ac PTM (Chen J. *et al*., 2013). All other combinations with H3K9me3 have negative interplay scores (Fig EV4). H3K9me3 therefore acts autonomously, independent of association with any other acetylation or methyl PTMs and not influenced by *P. falciparum* life cycle development. This feature of H3K9me3 is supported in other cell types where H3K9me3 act to reprogram the identify of various cell types (Becker *et al*., 2016; Nicetto & Zaret, 2019).

Several PTMs (45) display positive interplay throughout development, implying coordinated function or co-dependence including H3R17me2, H3K14me1 and H3R40me1 found in various combinations with partner PTMs, particularly H3K4me2/3 (Fig 4A, Fig EV4). These marks therefore likely never function on their own and needs interaction with one (or more) PTMs throughout parasite development. Interestingly, a number of these marks show increased positive interplay scores in mature gametocytes, suggesting that these combinations are increasingly critical these parasites. H3K14ac remains co-dependent on H3K9ac as seen before for asexual parasites (Fan *et al*., 2004a, b) with pronounced dependency evident in gametocytes here (Fig 4); indeed, the effector protein, GCN5 is expressed in all three stages investigated including mature gametocytes (Lopez-Barragan *et al*., 2011).

A set of 12 combinations are found in all three life cycle stages, yet have a diametrically opposed profile between gametocytes (strong positive interplay scores) and asexual parasites (strong negative interplay) (Fig 4B, Fig EV4). This includes a number of combinations involving H3K9me2, H3K36me1 and H3K18me1. The repressive mark H3K9me2 associates with several other marks (e.g. R17me1, K18me1, K36me1, K37me1 and R40me1), all of which show co-dependency only in mature gametocytes. The strongest differential interplay score between asexual and gametocytes was observed for H3K4me3K23me1. This novel combination is mutually exclusive in trophozoites, supporting the fact that H3K4me3 participates in crosstalk with H3K9ac in asexual parasites (Salcedo-Amaya *et al*., 2009). The strong co-dependence of H3K4me3K23me1 is therefore likely important for gametocyte-specific biological processes; the combination of the euchromatic H3K4me3 with H3K23me1 methylation has been associated with heterochromatin in *C. elegans* (Vandamme *et al*., 2015).

The conserved nature of these co-existing marks across all life cycle stages of *P. falciparum* imply shared importance to parasite biology.

### A dynamic histone code describes crosstalk with stage-specific biology in P. falciparum

Several PTMs show a crosstalk profile that is uniquely associated with a specific parasite life cycle stage (Fig 5). In trophozoites, the majority of the PTM combinations had negative interplay scores, suggesting that these PTMs antagonize each other, and are thus mutually exclusive. Combinations including novel arginine methylation marks like H3K14me2R17me1, show the strongest negative interplay scores (IS=-3.7), suggesting that these marks act with different biological roles from each other in trophozoites (Fig 5A). In fact, both H3R40me1 and H3R17me1 mostly show pronounced negative interplay scores for the majority of their interactions in trophozoites. Whilst H3R17me1 is activating as de-repressor in mammalian cells (Miller & Grant, 2013), H3R40me1 has only previously been reported in a few organisms including yeast where it required for efficient sporulation, similar to spermatogenesis in higher eukaryotes (Govin *et al*., 2010; Huang & Hull, 2017), while in cancer cells (Li Q.Q. *et al*., 2017), *C. elegans* (Sidoli *et al*., 2016b) and murine embryonic stem cells, its function is unknown (Sidoli *et al*., 2014).

**Figure 5.**
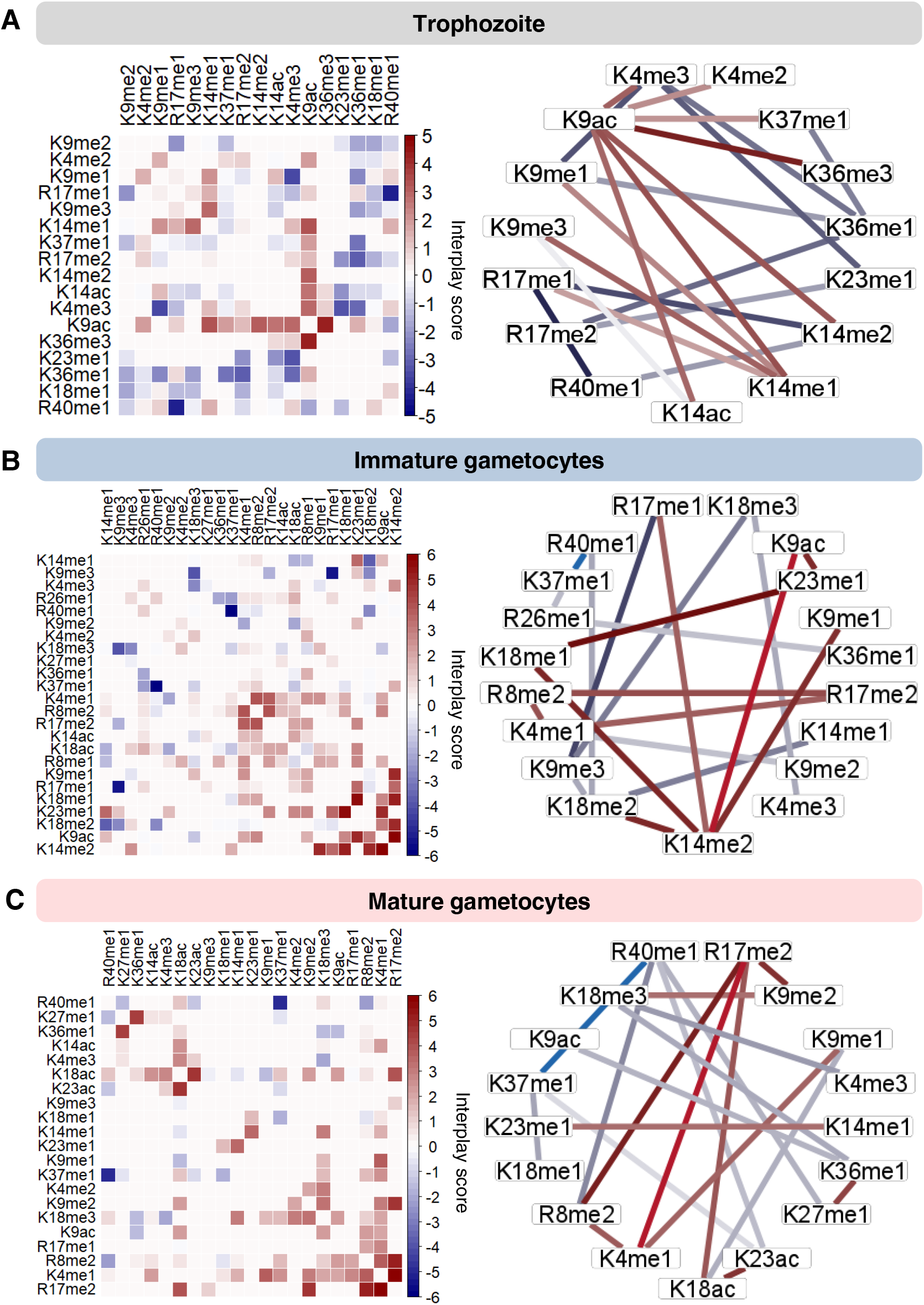
Dynamic, stage-stratified histone modification crosstalk in *P. falciparum* parasites. Interplay scores between bivalent PTMs occurring in **(A)** trophozoites, **(B)** immature gametocytes and **(C)** mature stage gametocytes where the colour intensity of the square is proportional to the interplay score. Ring plots indicate top and bottom 10 bivalent combinations, where the colour of the edges are proportional to the interplay score shown in the heatmap. Interplay scores for trophozoites include the unique 4 % and the overlapping 6 % with immature gametocytes (I_GC) and 1 % with mature gametocytes (M_GC). Interplay scores for immature gametocytes include the unique 25 % and the overlapping 6 % with trophozoites and 25 % with mature gametocytes. Interplay scores for mature gametocytes include the unique 6% and the overlapping 4 % with trophozoites and 25 % with immature gametocytes.

Several important co-dependent interactions with positive interplay scores were identified in trophozoites, the majority of which involved H3K9ac (e.g. H3K9acK14ac IS = 2.8; H3K4me3K9ac IS= 2.9; and H3K9acK36me3) to produce the global euchromatic signal associated with asexual parasites (Bartfai *et al*., 2010; Salcedo-Amaya *et al*., 2009). H3K9acK36me3 has the strongest positive interplay score (IS=4.3) and both H3K9ac and H3K36me3 have been independently linked to *var* gene transcription (Bartfai *et al*., 2010; Connacher *et al*., 2021; Jiang L. *et al*., 2013; Karmodiya *et al*., 2015; Rando, 2007); this co-dependence supports their function in *var* gene expression.

The crosstalk profile in immature gametocytes is more complex, with a larger number of PTMs involved in positive crosstalk, the majority of which involve methylation marks (Fig 5B). The novel mark H3K14me2 displays the largest number of interactions with positive crosstalk, including that with H3K9ac (IS=6), H3K9me1 (IS=4.9) and with both K18me1&2 (IS∼5). These associations are influenced by the level of methylation (i.e. mono-, di- or trimethylation) to define chromatin states and active genes *vs.* inactive genes within the same locus (Karachentsev *et al*., 2007; Schneider *et al*., 2004; Wang *et al*., 2018), with H3K14me1 for instance in negative crosstalk with H3K18me2, compared to the positive crosstalk seen for H3K14me2 with H3K18me2. H3K14me2 is likely a repressive mark, similar to the important silencing function of H3K14me3 that mark a set of zinc finger protein genes during trans-differentiation of bone marrow cells into hepatocytes (Liao *et al*., 2015; Zhao B. *et al*., 2018). The co-dependency with H3K14me2K9me1 (with H3K9me1 as key PTM in the establishment of functional heterochromatin (Grewal & Rice, 2004) and the novel mark H3K14me2K18me1/2, marks H3K14me2 as a key modulator of gene regulation within immature gametocytes, to potentially mediate the establishment of a heterochromatic state.

Amongst the novel arginine PTMs in immature gametocytes, H3R17me2 show positive interplay particularly with H3R8me2 (IS = 4.1, correlated only with active histone PTMs (Dong *et al*., 2018) and H3K4me1 (IS = 3.6, enhancer associated activation mark (Bae & Lesch, 2020), with the latter also positively connected (H3K4me1R8me2, IS = 3.97). H3R17me2 is a typical activation mark in eukaryotes (Di Lorenzo & Bedford, 2011; Vanagas *et al*., 2020; Wu & Xu, 2012) and the coordination with the two other activation marks indicate that these combinations may be involved in specific activation processes. Interestingly, changing H3R17 methylation from di- to monomethylation, results in positive interactions with H3K14me2 (IS = 3.59) but also a strong negative interaction with H3K9me3 (IS = - 5.33), implying independence of H3R17me1 from H3K9me3. Furthermore, in immature gametocytes, H3R40me1 again show strong negative interplays, particularly with H3K37me1 (IS = -7) and H3K18me2 (IS= -2.8), confirming its independent action in immatures gametocytes, similar to in trophozoites. Overall, immature gametocytes use highly connected and co-dependant repressive lysine PTMs pairs to induce a more heterochromatic state compared to asexual parasites. However, the majority of arginine PTM coordinate positively and could result in activation of subsets of genes.

Mature gametocytes have a combinatorial profile similar to immature gametocytes with largely positive interplays scores (Fig 5C), although the frequency of the combinations somewhat differ from those in immature gametocytes. H3R40me1 remains highly connected in mature gametocytes, again showing independence from other co-existing marks. Activating H3R17me2 is most connected PTM in mature gametocytes, retaining its interactions and co-dependency with other activating marks, H3K18ac (IS = 3.9) and H3K4me1 (IS = 5.9), and also with H3K9me2 (IS = 4.8) and H3R8me2 (IS = 5.2). Mature gametocytes are, however, additionally marked by combinations that uniquely and exclusively is present only in this stage of development. This includes co-dependency between H3K27me1K36me1 (IS = 4.3). In embryonic stem cells, H3K27me1 is dependent on H3K36me1 to promote transcription (Jung *et al*., 2013) and an increase in H3K36 methylation to di- and trimethylation negatively impacts H3K27 function and they become mutually exclusive (Ferrari *et al*., 2014; Zheng *et al*., 2012). H3K36me2&3 is repressive to asexual gene sets in immature gametocytes (Connacher *et al*., 2021). The exclusive combination of H3K27me1K36me1 in mature gametocytes could therefore be predicative of transcriptional activation of genes required for parasite transmission. Additionally, H3K18ac also is in crosstalk with H3K23ac only in mature gametocytes (IS = 4.7 for H3K18acK23ac). H3K18ac independently associates to active promotors in asexual *P. falciparum* parasites (Tang *et al*., 2020) and in human cancer cells, deacetylation of H3K18ac by SIRT7 results in transcriptional repression (Barber *et al*., 2012). Furthermore, the co-dependency between H3K18ac, H3K23ac and H3R17me2 have been demonstrated to induce transcriptional activation in cancer cells (Daujat *et al*., 2002) and could therefore similarly be critical for stage-specific gene expression exclusively in mature gametocytes. This coordination may be an essential requirement for subsequent gamete formation and fertilisation. The histone code in mature gametocytes may therefore result in bistable chromatin to enable a transcriptionally poised state of some genes for rapid fertilisation and sexual replication once these mature gametocytes are taken up by a feeding mosquito, and this is mediated by unique co-dependent histone PTM combinations.

### Shared protein effectors coordinate to mediate H3K18acK23ac function in mature gametocytes

The unique combination of H3K18acK23ac, and the associated co-existence with H3R17me2 in mature gametocytes, warranted further investigation of the presence of shared effector proteins to promote a biological outcome of transcriptional activation associated with these combinations. To identify the protein machinery associate with these marks, we performed quantitative chromatin immunoprecipitation on crosslinked chromatin isolated from mature gametocytes, coupled with quantitative mass spectrometry (ChIP-MS, Fig 6A). The presence of both marks found in combination was validated by western blot analysis in mature gametocytes (Fig EV5A). Since an antibody against H3K18acK23ac is not commercially available, we opted to capture proteins that interact with or are recruited to both H3K18ac and H3K23ac, respectively, in two separate ChIP experiments using antibodies specific to H3K18ac and H3K23ac (Fig EV5B). We used this technique to ensure that any endogenous protein (or protein complexes) in mature gametocytes that associate with these marks is captured in its *in vivo* context (Wierer & Mann, 2016).

**Figure 6.**
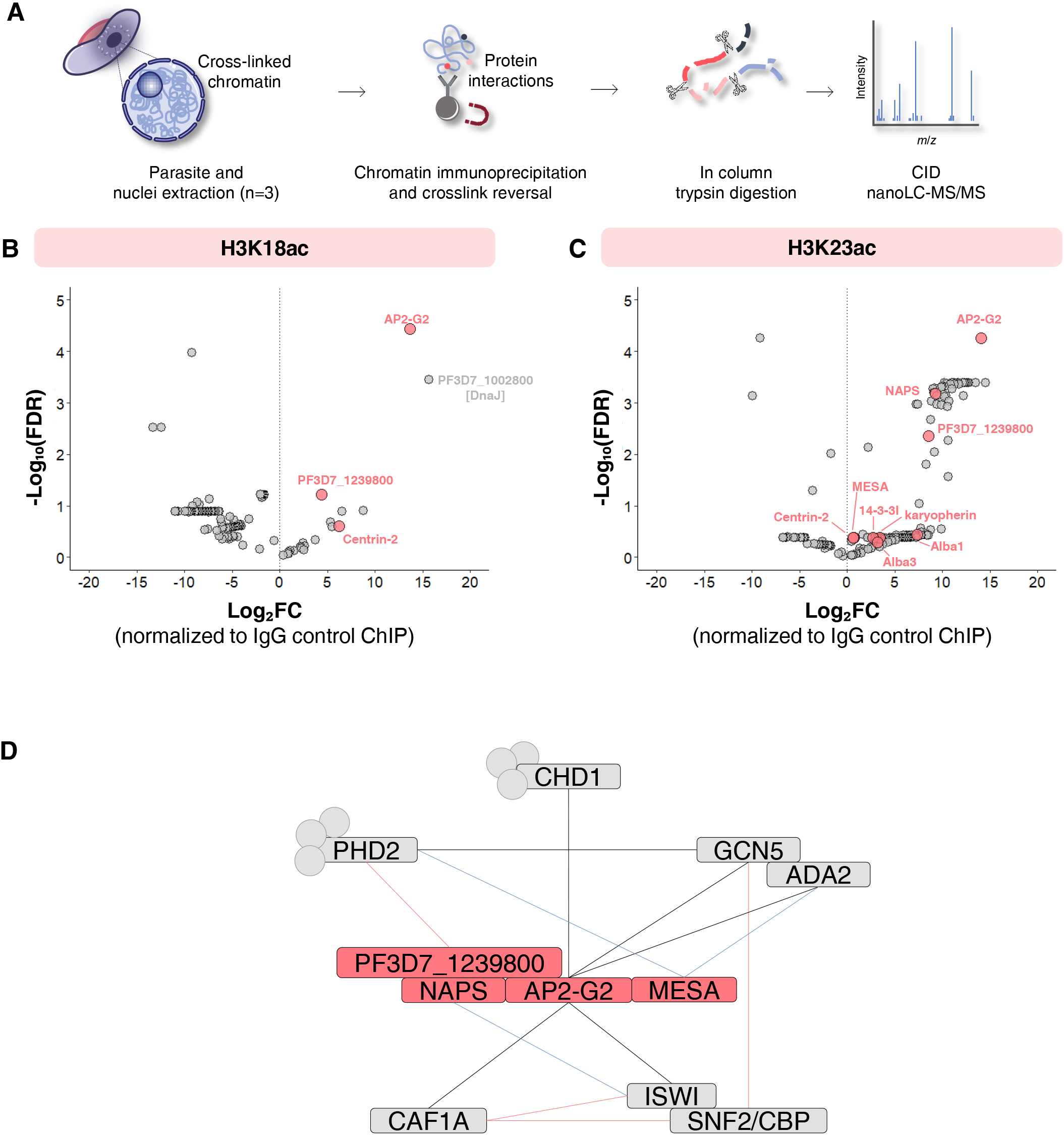
Proteins identified by chromatin proteomic profiling that are associated with H3K18ac and H3K23ac in mature stage gametocytes. **(A)** After parasite DNA and proteins were crosslinked, nuclei were isolated, and chromatin sonicated. Chromatin complexes were immunoprecipitated with antibodies raised against histone PTMs H3K18ac and H3K23ac. Crosslinks were reversed, and proteins trypsin digested. Peptides were fractionated with CID for MS analysis. NanoLC-MS/MS was performed, followed by database searching using Thermo Proteome Discoverer (v.1.4.1.14) to extract peaks, Scaffold™ (v.4.5.3) to validate and quantify peptides, Mascot (v.2.5.1) to identify proteins, and proteins identified searches using databases PlasmoDB (v.46) and UniProt (2020_01). Proteins were quantified using iBAQ values. Images were adapted from Servier Medical Art (URL link to the license: https://creativecommons.org/licenses/by/3.0/) and changes were made in terms of colour, size and composition. Scatter plot of the **(B)** H3K18ac and **(C)** H3K23ac-associated proteome enrichment in the ChIP-MS. A total number of 282 proteins were identified in the preparations, with several proteins showing positive log₂-fold change normalized to the negative control IgG ChIP. Proteins that are shared between the H3K18ac and H3K23ac samples include transcription factor AP2-G2 (PF3D7_1408200), a conserved unknown function protein (PF3D7_1239800) and centrin-2 (PF3D7_1446600) as shown in pink. H3K23ac also includes nucleosome assembly protein (PF3D7_0919000, NAPS), DNA/RNA-binding protein Alba 1 (PF3D7_0814200, Alba1), karyopherin beta (PF3D7_0524000), DNA/RNA-binding protein Alba 3 (PF3D7_1006200), 14-3-3 protein (PF3D7_0818200, 14-3-3I), mature parasite-infected erythrocyte surface antigen (PF3D7_0500800, MESA). **(D)** A schematic model of the protein-protein interaction complex manually curated and placed in a network based on evidence identified in this study, a AP2-G2 peptide pulldown study (Singh *et al*., 2020) and a GCN5 pulldown (Hoeijmakers *et al*., 2019), previous yeast two hybrid protein interactions (La Count *et al*., 2005) and STRING interactions. Shown in grey blocks are interactions that include the transcriptional coactivator ADA2 (PF3D7_1014600, ADA2), histone acetyltransferase GCN5 (PF3D7_0823300, GCN5), chromodomain-helicase-DNA-binding protein 1 homolog (PF3D7_1023900, CHD1), ISWI chromatin-remodelling complex ATPase (PF3D7_0624600, ISWI), Snf2-related CBP activator (PF3D7_0820000, SNF/CBP), chromatin assembly factor 1 subunit A (PF3D7_0501800, CAF1A), and PHD finger protein PHD2 (PF3D7_1433400, PHD2). Grey dots represent other proteins that associate to the respective proteins but are inconsequential. Proteins shown in pink blocks are shared between the H3K18ac and H3K23ac samples include transcription factor AP2-G2 (PF3D7_1408200), nucleosome assembly protein (PF3D7_0919000, NAPS), a conserved unknown function protein (PF3D7_1239800) and mature parasite-infected erythrocyte surface antigen (PF3D7_0500800, MESA). Black edges represent data from the AP2-G2 and GCN5 interactomes (Singh *et al*., 2020 and Hoeijmakers *et al*., 2019); the pink line represent data from STRING [(STRING: functional protein association networks (string-db.org)]; and the blue line indicated data from a yeast to hybrid study (La Count *et al*., 2005).

We defined proteins significantly associated with H3K18ac and H3K23ac as those with a log_2_ fold change ≥ 2 for the ChIP population (crosslinked chromatin preparation) compared to the IgG control ChIP, at a FDR ≤ 5%. Selective enrichment of proteins for H3K18ac and H3K23ac was confirmed by: 1) the significant (*P* ≤ 0.01) enrichment of unique peptides for low abundance proteins in the H3K18ac and H3K23ac ChIP preparations compared to the IgG sample preparations (Fig EV5C); 2) significantly increased abundance of proteins in the histone PTM samples compared to the IgG samples based on relative intensity-based absolute quantification (iBAQ) (Fig EV5D, *P* ≤ 0.01) and lastly, 3) the enrichment in the ChIP population for chromatin associated proteins (>20% enrichment)(Batugedara *et al*., 2020), the nuclear pore proteome (Oehring *et al*., 2012), and transcription factor interacting proteins (Singh *et al*., 2020)(Fig EV5E).

We found three proteins to be strongly enriched for H3K18ac in the ChIP-MS data, compared to 46 proteins for H3K23ac (FDR < 5%, log_2_ fold change ≥2 over IgG control ChIP)(Fig 6). The transcription factor AP2-G2 (PF3D7_1408200), a protein with an unknown biological function (PF3D7_1239800), centrin-2 (PF3D7_1446600) and DnaJ (PF3D7_1002800) were enriched in both PTM ChIPs. Besides, the chaperone DnaJ, all of these proteins were previously implicated to be chromatin associated (Batugedara *et al*., 2020). The gene for PF3D7_1239800 is refractory to deletion, is likely essential for asexual parasite survival (Zhang *et al*., 2018) and is highly expressed in mature gametocytes (Lopez-Barragan *et al*., 2011). AP2-G2 is also essential for the successful completion of gametocyte maturation and transmission (Singh *et al*., 2020; Xu *et al*., 2020).

To further investigate the epigenetic complexes associated with H3K18ac and H3K23ac, protein-protein interaction data were used to assemble a proposed reader complex for these PTMs in mature stage gametocytes, associated with AP2-G2 (Fig 6C)(Hoeijmakers *et al*., 2019; LaCount *et al*., 2005; Singh *et al*., 2020) (Appendix 3). Transcription factors are indeed observed in many histone PTM-associated complexes (Hoeijmakers *et al*., 2019) to recruit additional members of epigenetic complexes, as shown for AP2-I (Santos *et al*., 2017). The complex included evidence for the direct interaction between GCN5 (PF3D7_0823300), with acetyltransferase activity, and ADA2 (PF3D7_1014600), pointing towards the involvement of a HAT SAGA-like complex. This was supported by AP2-G2 further interacting with the chromodomain-helicase-DNA-binding protein 1 homolog (PF3D7_1023900; CHD1), which in turns interacts with PHD2 (PF3D7_1008100), a parasite-specific PHD-finger domain containing protein with different specificity to its PHD1 partner (unable to bind e.g. H3K4me3, (Hoeijmakers *et al*., 2019)). The nucleosome assembly protein (NAPS) and PF3D7_1239800 also has direct interactions with PHD2. In gametocytes, H3K18acK23ac therefore associates with a GCN5-ADA2-PHD2 SAGA-like complex via the tight interaction of AP2-G2, NAPS and PF3D7_1239800 as binding partners to this particular histone combination. Most of these proteins are predominantly expressed in male mature gametocytes compared to female gametocytes, including ADA2, PHD2, ISWI, NAPS, CHD1 and AP2-G2 (Lasonder *et al*., 2016), implicating these proteins in downstream sex-specific chromatin structure changes.

AP2-G2 additionally interacts with other chromatin regulation proteins, including a chromatin assembly factor 1 subunit A (PF3D7_0501800, CAF1A) and a ISWI chromatin-remodelling complex ATPase (PF3D7_0624600, ISWI), which interacts with a Snf2-related CBP activator (PF3D7_0820000). Both ISWI and Snf2/CBP are homologues of the mammalian H3K18acK23ac writer and reader, p300 and TRIM24, respectively (Halasa *et al*., 2019; Luo *et al*., 2019; Lv *et al*., 2017; Ma *et al*., 2016; Tsai *et al*., 2010). This suggest that the SAGA-like complex of *P. falciparum* has as core GCN5/ADA2/PHD2 but in gametocytes, association with the H3K18acK23ac combination includes the additional ISWI/SNF complex effectors. This cooperation between SAGA and SWI/SNF complexes is required to regulate specific transcriptional responses, as in yeast (Sanz *et al*., 2016). Since SWI/SNF complex proteins are global nucleosomal organizers that enable the specific binding of selective transcription factors (Barisic *et al*., 2019; Dutta *et al*., 2017; Mohrmann *et al*., 2004), their involvement could explain the recruitment of AP2-G2.

The involvement of Snf2/CBP further points to functionality for the crosstalk involving the neighbouring H3R17 mark, forming the H3R17me2K18acK23ac coordinating code in *P. falciparum* mature gametocytes. Arginine 17 methylation is achieved in a systematic manner: CBP first acetylates H3K18, then H3K23 and this allows the arginine methylase CARM1 to associate with chromatin to methylate H3R17 (Sakabe & Hart, 2010; Yue *et al*., 2007). H3R17me2 is associated with transcriptional activation based on the recruitment of polymerase-associated factor 1 complex to initiate transcription in humans (Shishkova *et al*., 2017; Wu & Xu, 2012), supported by the antiproliferative effects of a specific CARM1 inhibitor on multiple myeloma cell lines (Drew *et al*., 2017; Li Y. & Seto, 2016). This suggest that in *P. falciparum*, this combination could be critical for mature stage gametocytes.

Given the shared core proteins between H3K18ac and H3K23ac and the positive crosstalk observed for this combination (and including R17me2), our data for this combination provides evidence that the combinatorial histone code of the *P. falciparum* parasite can recruit protein complexes unique to a combination to facilitate downstream biological processes.

## Discussion

Here we present the first systems-level evidence of a comprehensive combinatorial histone code for various life cycle stages of the human malaria parasite, *P. falciparum*. Middle-down proteomics provided high-resolution quantitative data to describe the histone code in this parasite, which could function as model for other protista. Our data reveal that the combinatorial histone PTM landscape is dynamic, with clear stage-specific differentiation observed, with gametocyte stages more dependent on histone PTM crosstalk than asexual parasites.

Collectively, our study shed light on the difference between the asexual replicating parasite and the differentiated non-replicative gametocyte. The histone code of *P. falciparum* asexual parasites resembles that of lower eukaryotes, which has a simple genome organisation and fewer histone PTMs. Given the primal function it performs, elaborate epigenetic gene regulation mechanisms may not be as important to these stages, as most of the genome is in an euchromatic state and actively transcribed during proliferation. Therefore, asexual parasites use less, and mutually exclusive, histone PTMs to regulate key functions such as host immune evasion. However, gametocytes share a similar mechanism of epigenetic gene regulation with other higher-order, multicellular, eukaryotes where chromatin is predominantly condensed and highly regulated to specify the identity and purpose of a cell through multiple histone PTMs. The majority of gametocytes indicate a general positive crosstalk between histone PTMs and given the limited number of transcription factors in *P. falciparum* parasites, gametocytes could rather switch to a more complex epigenetic code to impress very specific regulation of its biological processes. This would suggest that at least in some stages of the parasite, epigenetic level gene control is superior to transcriptional level control.

The connectivity within the histone code of *P. falciparum* is characterised by the presence of novel marks, with several new arginine modifications identified. The advances in proteomics technologies such as middle-down proteomics is allowing such robust description of arginine methylation marks (Li Q.Q. *et al*., 2017) as observed here, and this contributes to our understanding of the conserved nature of arginine methylation and its key importance to chromatin organisation throughout eukaryotes (Di Lorenzo & Bedford, 2011). We show that histone arginine methylation is equally as prevalent and abundant as lysine PTMs in *P. falciparum* and these marks participate in complex crosstalk with one another, particularly in gametocytes. The presence of typically activating marks such as mono- and demethylation of H3R17 (Di Lorenzo & Bedford, 2011) and their co-dependence on other marks e.g. H3R8me2 raise interesting questions as to the importance of cooperation in activation of gene sets in gametocytes. This is extended to additional arginine methylation marks including the exclusive nature of H3R42me1 in gametocytes and the highly connected, but independently functioning H3R40me1. It is noteworthy that H3R40me1 is required as activating mark for spermatogenesis-like processes in yeast (Govin *et al*., 2010; Huang & Hull, 2017). These marks could be similarly important for activation of male gamete gene sets, supporting the notion of transcriptionally active ‘poised’ states in mature gametocytes (van Biljon *et al*., 2019) to enable onwards transmission. The importance of arginine methylation in histone PTM combinations to mediate a specific transcriptional outcome in gametocytes is therefore of interest. The parasite genome does contain the necessary machinery for arginine methylation including two putative protein arginine methyltransferases (PRMT1 [PF3D7_1426200] and PRMT5 [PF3D7_1361000]) and a putative histone arginine methyltransferase (CARM1/PRMT4 [PF3D7_0811500]). Indeed, PRMT inhibitors are active against *P. falciparum* parasites (Fan *et al*., 2009) and CARM1/PRMT4 is essential in asexual parasites (Zhang *et al*., 2018), supporting functional importance of these marks. Protein arginine methyltransferase inhibitors are seen as promising anticancer targets (Hwang *et al*., 2021) and could be applied in the malaria context for gametocyte-targeting, transmission-blocking compounds.

The connectivity of histone PTMs in the *P. falciparum* histone code is clearly associated with different developmental outcomes, similar to other eukaryotic systems requiring specialisation e.g., embryogenesis and stem cell differentiation (Atlasi & Stunnenberg, 2017; Vastenhouw & Schier, 2012). Importantly, the prevailing heterochromatic mark, H3K9me3, is highly connected but not co-dependent on any other acetylation or methylation mark across all the life cycle stages. In the absence of quantified association between H3S10ph, as co-existing mark impairing HP1 binding to H3K9me3 (Fischle *et al*., 2005) in Plasmodia, this indicates that H3K9me3 is likely singularly important to binding of HP1 to demarcate heterochromatin in *P. falciparum*. However, gametocytes additionally use other highly connected, but independently acting marks repressive marks such as H3R40me1 and H3K14me2 to likely govern strategy-specific gene inactivation, as has been described for H3K36me2/3 inactivation of gene sets typically only required in asexual parasites (Connacher *et al*., 2021). The observation that repressive marks are highly connected but in the majority of instances show independence or mutual exclusivity to the partner PTMs questions the importance of repression in Plasmodia. The limited set of histone PTMs to enable effective transcriptional repression and induction of heterochromatin could be important to control transcript levels of particular gene sets, e.g. virulence genes in asexual parasites (Jiang L. *et al*., 2013) but are more important during gametocytogenesis. Additional mechanisms including RNA decay may be more influential to regulate transcript levels in general (Painter *et al*., 2017; Shock *et al*., 2007).

Connectivity between PTMs in *P. falciparum* is evidently important for coordinated function and activation of euchromatin. Silencing PTMs usually form independent heterochromatic domains (e.g. H3K9me3), but activating marks are frequently found together. The majority of activating PTMs coordinate irrespective of life cycle stages, particularly H3K9acK14ac and explains the euchromatic permissive nature of asexual parasites, enabled by coordinated binding of effector proteins. Novel combinations such as H3K4me3K23me1 and several combinations with H3R17me1 and H3R17me2 (H3K4me1, H3R8me2, H3K9me2) is pronounced during gametocytogenesis. Additionally, during these stages of unique differentiation of *P. falciparum*, the parasite relies on specific and differentiated combinations, including the unique H3K27me1K36me1 and H3R17me2K18acK23ac combinations seen in only mature gametocytes. The connectivity and crosstalk between histone PTMs are therefore essentially important to establish general euchromatic regions in the parasite’s genome across all stages. However, crosstalk of activating marks is more prevalent in gametocytes and requires differentiation in marks used for euchromatin in these stages. This implies the use of specific histone combinations for activation of strategy-specific gene sets to mediate *P. falciparum* transmission.

The functional relevance of the connectivity of histone PTMS is underscored by evidence that the associated interacting proteins are also highly connected to include reader and writer proteins, ‘flavoured’ to specific marks (Hoeijmakers *et al*., 2019). In this manner, the crosstalk in the PTM combinations results in recruitment of protein complexes to interpret the PTM combinations and allow changes to the chromatin structure. Indeed, we show that the unique co-dependent combination in mature gametocytes, H3K18acK23ac, jointly recruits the transcription factor AP2-G2, to initiate a SAGA-like complex containing GCN5/ADA2, flavoured with K18acK23ac-specific effectors. Since co-dependency extends to the triple combination H3R17me2K18acK23ac, we provide evidence that this combination may indeed functionally associate with a gametocyte-specific SAGA-like complex to mediate stage-specific gene expression exclusively in mature gametocytes, as this combination have been demonstrated to induce transcriptional activation in cancer cells (Daujat *et al*., 2002). The identification of AP2-G2 as immediate binding partner on the H3K18acK23ac combination suggests that in mature gametocytes, AP2-G2 may act as transcriptional activator by targeting these histone PTM combinations. Investigations of the specific gene sets controlled by H3R17me2K18acK23ac and AP2-G2 in mature gametocytes are underway.

The histone code in *P. falciparum* is therefore diverse and dynamic to effect different combinations required for proliferation and differentiation. The complex nature of the combinations and changes in the identity of the combinations during stage transition points to this epigenetic level of regulation being a more important level of regulation that can be easily finetuned by variation in the combinations, particularly for gametocytes. With a limited set of effector proteins and only a core of ∼5 full effector protein complexes characterised (Hoeijmakers *et al*., 2019), the intricacy of the histone code indicates that indeed, combinations of histone PTMs provides the blueprint to ensure differentiation, specificity and variation to control transcriptional activation of gene sets during *P. falciparum* development.

In conclusion, our study contributes a comprehensive catalogue of histone PTM combinations and provide a foundation for further investigation of the increasingly intricate histone code of the *P. falciparum* parasite.

## Materials and Methods

### *In vitro* cultivation of *P. falciparum* asexual parasites and gametocytes

All *in vitro* experiments involving human blood donors and human malaria parasites holds ethics approval from the University of Pretoria Research Ethics Committee, Health Sciences Faculty (NAS332/2019). Intra-erythrocytic *P. falciparum* parasites (NF54 strain, drug sensitive) was cultivated in fresh human erythrocytes (either A⁺ or O+) in RPMI-1640 culture medium supplemented with 25 mM HEPES (pH 7.5, Sigma Aldrich, USA), 0.2 mM hypoxanthine (Sigma Aldrich, USA), 0.024 µg/µL gentamycin (Hyclone, USA), 5 µg/µL Albumax II (Invitrogen, USA), 23.81 mM sodium bicarbonate (Sigma Aldrich, USA) and 0.2 % w/v D-glucose. Cultures was maintained with daily media change and fresh erythrocyte supplementation at 5 % haematocrit, 2 % parasitaemia under hypoxic conditions (5 % O₂, 5 % CO₂, 90 % N₂) with moderate shaking at 37°C. Parasites were synchronised to more than 90 % rings stages with D-sorbitol. Gametocytogenesis production was initiated at 0.5% parasitaemia and at a 6 % haematocrit in a glucose-free medium under hypoxic gaseous (5 % O₂, 5 % CO₂, 90 % N₂) conditions at 37°C without shaking (Reader *et al*., 2015).

### Histone isolation

Nuclei were liberated from *P. falciparum* parasites using a hypotonic buffer containing 10 mM Tris-HCl (pH 8.0), 3 mM MgCl₂, 0.2 % v/v Nonidet P40, 0.25 M sucrose and a cocktail of EDTA-free cocktail of protease inhibitors. Histones were subsequently extracted from chromatin and enriched through incubation with of 0.25 M hydrochloric acid. Histones were precipitated using 20 % trichloroacetic acid and rinsed with acetone air-dried and reconstituted using dddH_2_O. Histones were subsequently concentrated by freeze drying and stored at -80°C.

### Middle-down proteomics for identification of combinatorial histone PTM

Middle-down MS was performed according to Coradin *et al*. (2020) with a few modifications (Coradin *et al*., 2020; Sidoli *et al*., 2016b; Sweredoski *et al*., 2017). Histones (15-50 µg) were suspended in 5 mM ammonium acetate to a final concentration of 0.5 µg/µL (pH 4.0). GluC endoprotease was diluted to 0.2 µg/µL in the same buffer and added to the histone samples a final concentration of 1:20 GluC enzyme:histone. Histone samples were incubated for overnight at room temperature and digestion is blocked by addition of 1 % formic acid. Trifluoroacetic acid (0.1 %) was added to the digested peptides and desalted using StageTips packed with a solid phase C_18_ column disk (3M Empore) with porous graphitic carbon (PGC) suspended in 100 % acetonitrile (ACN) on top and washed with 0.1 % trifluoroacetic acid. Peptides were eluted by the addition of 70 % acetonitrile and 0.1 % trifluoroacetic acid elution buffer. Histone protein samples were dried and suspended to a concentration of ∼2 µg/µL in the sample buffer consisting of 60 % acetonitrile, 20 mM propionic acid and ethylenediamine.

Separation of digested histone peptides was performed by using nanoliter flow liquid chromatography using an EASY-nLC nanoHPLC (Thermo Scientific, San Jose, CA, USA) equipped with an analytical weak cation exchange–hydrophilic interaction liquid chromatographic resin (PolycatA, PolyLC, Columbia, MD, USA) column (75 μm ID, 15 cm in length, 1.7 μm in diameter and a 1000 Å porosity). Buffer A was prepared with 70 % acetonitrile and 20 mM propionic acid to adjust the pH to 6.0. Buffer B consisted of 0.1% formic acid in LC-MS grade H_2_O. HPLC gradient was setup for a nonlinear gradient: 0-70 % of buffer B for 2 min followed with 72-85 % buffer B for more than 90 min. The gradient is on hold for 5 min at 95 % buffer B to wash the column prior to consequential sample loading. To minimize sample carryover, blanks were run frequently for at least 20 min with high (80-95 %) buffer B. MS detection was performed using an Orbitrap Fusion mass spectrometer (Thermo Scientific, San Jose, CA, USA) operated in a data dependent mode and high mass resolution mode for both MS1 events and MS2 scans for each of the three stages investigated in two to three independent biological experiments. The full mass spectrum scan was set at 665–705 *m/z*, as this is the range of the most intense charge states (+8) for histone H3 polypeptides. A charge filter was added to include +8 charge states targeted for fragmentation. The ion transfer tube temperature was set to 300°C and the spray voltage to 2.3 kV. To fragment peptides while retaining PTM information, electron transfer dissociation (ETD) performed (Riley & Coon, 2018; Riley *et al*., 2017) at a resolution of 120 000 (MS1) and 30 000 (MS2). The ETD reaction time was 20 ms for polypeptides with +8 charge states. For high resolution spectrums, three microscans were averaged.

### Middle-down proteomics data analysis

Raw MS files were deconvoluted with Xtract (Thermo) and searched using Mascot (v. 2.5) against histone sequences obtained from PlasmoDB resource (https://plasmodb.org/, version 43, released 25 April 2019). Files were searched with the following dynamic modifications: Acetylation (K), phosphorylation (ST), mono and dimethylation (RK), and trimethylation (K). The mass tolerance was set to 2.1 Da for precursor ions and 0.01 Da for fragment ions. IsoScale (http://middle-down.github.io/Software/) was used to confidently identify and quantify modified peptides. Only *c*/*z* fragment ions were allowed. PTMs were only accepted if there were at least one site determining ions on both sides of the assigned PTM.

Interplay scores between bivalent histone PTMs were calculated according to the following equation:

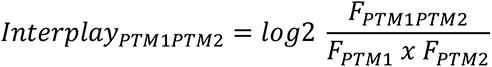

Where F_PTM1PTM2_ is the observed relative abundance (F) of the combination divided by the predicted relative abundance of the combination, calculated by multiplication of the relative abundances of the individual histone PTMs. The relative abundance of the individual PTMs were calculated by summing the relative abundances of all combinatorial peptides contain the particular PTM. If the interplay score is positive, it is likely that the PTMs are co-dependent or positively related; a negative interplay score indicates that the PTMs are mutually exclusive. Ring plots were visualised using Cytoscape (v. 3.8.2). Other plots were created using GraphPad Prism 9 and R (R Studio v. 4.0.3).

### Chromatin immunoprecipitation (ChIP)

A minimum number of isolated parasites (10^9^ cells/mL) was crosslinked with 1 % formaldehyde and subsequently quenched with 125 mM glycine. Crosslinked parasite nuclei were suspended in cold lysis buffer (10 mM HEPES pH 7.9, 10 mM KCl, 0.1 mM EDTA pH 8.0, 0.1 mM EGTA pH 8.0, 1 mM DTT and EDTA-free protease inhibitor cocktail) and transferred to a pre-chilled dounce homogeniser. NP40 was added to a final concentration of 0.25 % and parasites are subsequently lysed with dounce B for ∼100 strokes. Sonication shearing buffer (cocktail of protease inhibitors, 1 % SDS, 50 mM Tris-HCl (pH 8.0), 10 mM EDTA and 100 mM NaCl) were added to the nuclei pellet and sonicated with the BioRuptor UCD-200 (Diagenode, Belgium) for 25 cycles at high power and 30 s intervals. Crosslinks were reversed by incubating the input sample overnight at 65°C. The input sample was suspended to a final volume of 200 µL with ChIP dilution buffer (0.01 % SDS, 1 % Triton X-100, 1.2 mM EDTA, 16.7 mM Tris-HCl, pH 8.0, 150 mM NaCl and a cocktail of protease inhibitors). To dilute the SDS in the dilution buffer, Tris-EDTA buffer (1 M Tris, pH 8.0 and 0.5M EDTA, pH 8.0) was added with RNase to a final concentration of 0.2 µg/µL and incubated for 1 h at 37°C. The chromatin soup was incubated with 1 µg of either anti-H3K18ac or anti-H3K23ac, rotating at 4°C overnight. Protein G magnetic Dynabeads™ (Invitrogen by Thermo Fisher Scientific, Norway) was added and incubated for 2 h rotating at 4°C. A total 20 % of the eluted chromatin was then retained as the input. The remaining eluted material was then used for the western blot validation of combination histone PTM. the bead-chromatin complex was washed extensively with each of the following buffers with aspiration of the preceding buffer before addition of the next buffer: Low salt immune complex wash buffer (0.1 % SDS, 1 % Triton X-100, 2 mM EDTA, 20 mM Tris-HCl pH 8.1, 150 mM NaCl), high salt immune complex wash buffer (0.1 % SDS, 1 % Triton X-100, 2 mM EDTA, 20 mM Tris-HCl pH 8.1, 500 mM NaCl), LiCl immune complex wash buffer (0.25 M LiCl, 1 % NP-40, 1 % deoxycolate, 1 mM EDTA, 10 mM Tris-HCl pH 8.1) and TE Buffer (10 mM Tris-HCl pH 8.0,1 mM EDTA pH 8.0). The beads were suspended in fresh ChIP elution buffer (0.1 M NaHCO3, 1 % SDS) and the supernatant collected.

### Protein sample preparation for mass spectrometry analysis

Briefly, protein samples were mixed with an equal volume of methanol:chloroform (3:1 ratio), followed by a brief vortex step and centrifugation at 4°C. Samples were washed twice with methanol followed by centrifugation at 17,000 x*g* for 3 min and complete removal of methanol:chloroform. The resulting precipitated proteins were dried at room temperature. All samples were dissolved in a solution containing 6 M urea, 2 M thiourea and 50 mM ammonium bicarbonate, pH 7-8. Samples were subjected to reduction, alkylation and digestion in preparation for MS as follows. Samples were incubated at room temperature for 1 h in 5 mM DTT to reduce disulphide bonds, cysteine residues alkylated in 50 mM iodoacetamide and subjected to Lys-C (1-1.5 μg) and trypsin (10 μg) digestion at room temperature for 3 h. Samples were then diluted with 50 mM ammonium bicarbonate to lower the urea and DTT concentrations in solution to prevent trypsin inactivation. The samples were sonicated (Sonic Dismembrator Model 100, Fischer Scientific, USA) for 3 cycles of 10 s continuous sonication (output 2) followed by 10 s resting on ice to dissolve the pellets completely. Cysteine residues were alkylated by incubation with 40 mM iodoacetamide. This was followed by overnight digestion with 10 μg trypsin at room temperature. The pH of the trypsin digested samples was adjusted with ammonium hydroxide to ∼pH 8-8.3 followed by centrifugation at 17,000 x*g* for 1 min. StageTip (STop And Go Extraction Tip) clean-up in combination with protein fractionation was performed for all the samples using a dual resin approach, where both Empore™ C18 disks (3 M, USA) and OligoTM R3 reversed-phase resins were used in combination. This was done to remove unwanted contaminants before MS and to fractionate the samples to ensure optimal protein detection. The column was equilibrated with 1 mM ammonium bicarbonate (pH 8.0), after which the supernatant of each sample was added to the dual resin StageTip and pushed through the column using a syringe (dropwise). All samples were dried down completely under vacuum (SpeedVac concentrator). In preparation for the MS run, 0.1 % formic acid was added to the samples followed by ultrasonic bath sonication for 10 min at 4°C. The samples were centrifuged at 17,000 x*g* for 10 min and the supernatant was loaded onto a Dionex™ -LC system (Thermo Fisher Scientific, USA), coupled online with a Q-Exactive HF mass spectrometer (Thermo Scientific, USA). Peptides were loaded into a picofrit 20 cm long fused silica capillary column (75 μm inner diameter) packed in-house with reversed-phase Repro-Sil Pur C18-AQ 3 μm resin. A gradient of 105 min was set for peptide elution from 2-28 % buffer B (100 % ACN/0.1 % formic acid), followed by a gradient from 28-80 % buffer B in 5 min and an isocratic 80 % B for 10 min. The flow rate was set at 300 nL/min. MS method was set up in a DDA mode. The full MS scan was performed at 70,000 resolutions [full width at half maximum (FWHM) at 200 m/z] in the m/z range 350-1200 and an AGC target of 106. Tandem MS (MS/MS) was performed at a resolution of 17,500 with a Higher Energy Collision Dissociation (HCD) collision energy set to 20, an AGC target of 5×104, a maximum injection time to 100 ms, a loop count of 12, an intensity threshold for signal selection at 104, including charge states 2-4, and a dynamic exclusion set to 45 s.

### Chromatin proteomic profiling data analysis

MS raw files were analysed by MaxQuant software version 1.5.2.8. MS/MS spectra were searched by the Andromeda search engine against the *P. falciparum* UniProt FASTA database. Mass accuracy was set at 4.5 ppm for precursor and 20 ppm for the product mass tolerance. Peptides were filtered for high confidence (FDR<1 %) using Fixed Value validator. Intensity-based absolute quantification (iBAQ) enabled for label-free quantification, where iBAQ values are calculated by a MaxQuant algorithm (sum of peak intensities of all peptides matching to a specific protein/number of theoretical peptides). Match between runs was enabled and set to a 1 min window. All samples were run in triplicate for three independent biological replicates. For data analysis, iBAQ values were log₂-transformed and normalised by subtracting to each value the average value of the respective sample. The peptide relative ratio was calculated using the total area under the extracted ion chromatograms of all peptides with the same amino acid sequence (including all its modified forms) as 100 %. For isobaric peptides, the relative ratio of two isobaric forms was estimated by averaging the ratio for each fragment ion with different mass between the two species. Next, extracted ion chromatography of those m/z ions with a mass tolerance of 10 ppm were performed. Statistical significance was assessed using a two-tails heteroscedastic t-test (*P*-value representation * = < 0.05, ** = < 0.005, *** = < 0.0005).

### Protein-protein interaction model

STRING v.10 database [string.org, (Szklarczyk *et al*., 2015)] parameters were set to include protein-protein interactions based on evidence, sourced from neighbourhood, experiments, databases, co-occurrence and co-expression. The minimum required interaction score was required to be of high confidence (0.7) with a maximum of 100 proteins interacting the first shell and 100 proteins in the second shell. Data from peptide pulldowns studies were pooled from Singh *et al*., 2020 and Hoeijmakers *et al*., 2019 and filtered only to include proteins that were significantly enriched in the respective studies. Finally, protein-protein interactions derived from yeast-two-hybrid study (La Count *et al*., 2005). Taken together, proteins were imported into Cytoscape (version 3.8.2). Proteins were further filtered to include only known chromatin associated proteins and proteins that have homology to known H3K18ac and H3K23ac associated proteins.

## Acknowledgements

This work was supported by the South African Research Chairs Initiative of the Department of Science and Innovation, administered through the South African National Research Foundation (UID 84627) to LMB. The UP ISMC acknowledges the South African Medical Research Council (SA MRC) as Collaborating Centre for Malaria Research. SS gratefully acknowledges the Leukemia Research Foundation (Hollis Brownstein New Investigator Research Grant), AFAR (Sagol Network GerOmics award), Deerfield (Xseed award) and the NIH Office of the Director (1S10OD030286-01). Funding from the National Institutes of Health (grants AI118891 and CA196539) to BAG are gratefully acknowledged.

## Author Contributions

LMB conceived the study. HvG, MC, MM and JR conducted experiments, and analysed data with SS for interpretation. HvG and LMB wrote the paper with inputs from the other authors. All co-authors approved the final version of the paper.

## Conflict of Interest

The authors declare that they have no conflict of interest.

## Data Availability

The middle-down proteomics data generated in this study have been deposited in the Chorus database (https://chorusproject.org) and are accessible through project number 1721. All raw files from the ChIP-MS data are freely available on https://chorusproject.org at the project no. 1730. The data analysis pipeline meets all MIAPE standards.

